# Segment2P: Parameter-free automated segmentation of cellular fluorescent signals

**DOI:** 10.1101/832188

**Authors:** Noah Dolev, Lior Pinkus, Michal Rivlin-Etzion

## Abstract

The availability of genetically modified calcium indicators has made calcium imaging of neural signaling accessible and widespread whereby recording hundreds or even thousands of cells simultaneously is commonplace. Immunocytochemistry also produces large images with a great number of antibody labeled cells. A major bottleneck towards fully harnessing these techniques is the delineation of the neural cell bodies. We designed an online robust cell segmentation algorithm based on deep learning which does not require installation or expertise. The robust segmentation is achieved by pre-processing images submitted to the site and running them through DeepLabv3 networks trained on human segmented micrographs. The algorithm does not entail any parameter tuning; can be further trained if necessary; is robust to cell types and microscopy techniques (from immunocytochemistry to single and multi-photon microscopy) and does not require image pre-processing.

## 1. Introduction

Identifying cell bodies in order to extract calcium transients, count cells, or measure morphology are common practices in neuroscience. Currently, these tasks require either manual annotation, complicated parameter optimization and/or arduous installations. Two well-established software to date, for example, are CMNF-e [2, 3] for extracting calcium transients (which has several dozen parameters) and U-NET for naive segmentation (which requires setting up Caffe on a Linux server) for segmentation [4].

The combination of advances in multi-photon imaging at relatively high frame rate with advances in creating sensitive genetically encoded calcium indicators has made monitoring the activity of large neuron populations feasible [16]. However, extracting the calcium transients from a large number of cells in the presence of measurement noise, neural processes in the background and overlap between cell bodies creates a bottleneck in evaluating the data. To solve this, we designed a simple web interface, which accepts fluorescent micrographs of arbitrary size and type containing cell bodies, creates an array of pre-processed copies and feeds them into two DeepLabV3 architectures [5]. The two networks segment the images with different sets of hyperparameters (model 1 and model 2), to increase robustness of the results. These masks are then combined into a single mask by summing, thresholding and watershedding.

To train these models, we generated a novel training dataset by including human segmented calcium imaging projections of our own two photon retinal recordings using the Figure 8 platform [6]. Segmentations were performed by laymen given simple instructions but each segmented region of interest (ROI) was accepted only when segmentors passed a threshold level of agreement between them (20% of segmentors needed to agree on a pixel value). We also eliminated the work of segmentors that failed to properly segment 80% of the cells in several test images (which were segmented manually by experts in calcium imaging). These ground truth samples were then used to finetune Tsai et al’s Mask-R-CNN model [7]. We then segmented a larger number of calcium imaging micrographs using the Mask R-CNN, manually corrected this first-pass segmentation and added these to our training dataset. This dataset is available upon request.

We tested Segment2p on retinal ganglion cells imaged on two different two-photon microscopy setups and immunocytochemical slices from a neighboring lab. In our hands, Segment2p delineated cell bodies about as well as human segmentors.

## 2. Algorithm Description

Two DeepLabV3 models [5] were trained using Amazon’s SageMaker platform on the aforementioned dataset. Semantic segmentation algorithms are often compared based on their performance on large and diverse image test sets such as PASCAL VOC 2012 [13]. We chose the encoder-decoder atrous convolution DeepLabV3 architecture since it outperformed other models in terms of intersection over union on several industry test sets at the time of development [5, 13]. The two models were trained separately and their parameters were set as follows:

**Table 1:** Model Hyperparameters

| Parameter | Model 1 | Model 2 |
| --- | --- | --- |
| backbone | resnet-101 | resnet-101 |
| crop_size | 412 | 384 |
| early_stopping | TRUE | TRUE |
| early_stopping_min_epochs | 25 | 25 |
| early_stopping_patience | 50 | 50 |
| epochs | 1000 | 250 |
| learning_rate | 0.00303705 | 0.00027659 |
| lr_scheduler | poly | poly |
| mini_batch_size | 35 | 40 |
| momentum | 0.61335965 | 0.68091031 |
| num_classes | 2 | 2 |
| num_training_samples | 576 | 576 |
| optimizer | adagrad | adagrad |
| use_pretrained_model | FALSE | FALSE |
| validation_mini_batch_size | 16 | 16 |
| weight_decay | 0.00015608 | 0.00014451 |

Model one was trained with an objective of maximizing pixel accuracy 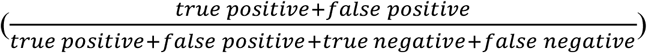 and model 2 was trained on an objective of maximizing the mean Intersection over Union (mIoU) (also known as the Jaccard index, see Section 3, Validating segment2p). Our algorithm reliably segments 2-photon micrograph projections of calcium imaging in spite of different dimensions [Fig 1]. Less than 1.67% of cells from the ground truth (e.g., cell bodies marked by human experts) are missed in the predicted segmentation shown in Figure 1 D. In addition, there were 5% more cells detected in the prediction than were delineated in the human segmented ground truth.

**Figure 1:**
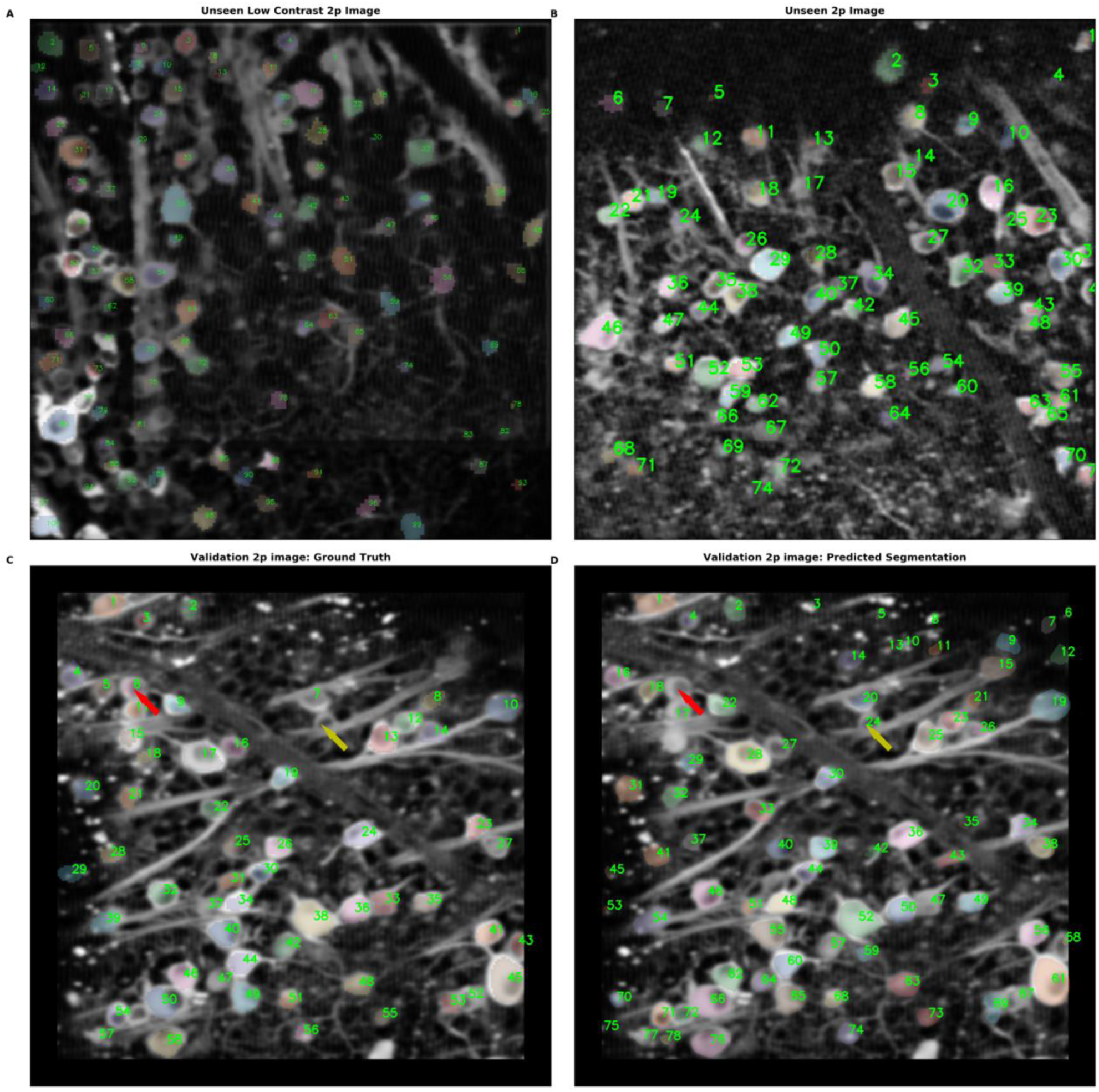
Sample segmentations of novel calcium imaging projections. Calcium image projections overlaid with segmentation masks. Top panel: Two never before seen 2p calcium image stack mean projections submitted to the website by students in our lab (data collected on different 2p microscopes with different protocols from training set). For presentation in the overlay, the images have been denoised via anisotropic diffusion and contrast enhanced (but submitted to segment2p without pre-processing). (A) Never before seen 1024×1024 pixel calcium image stack max projection. Cells appear low contrast compared to background but algorithm successfully performs segmentation. Note that the model correctly did not segment neural processes (B) Never before seen 256×256 calcium image stack max projection after contrast enhancement. (C) Sample validation image overlaid with the ground truth. (D) Same sample validation image overlaid with predicted segmentation. Red arrow indicates ROI present in ground truth but not predicted by model and yellow arrow represents cell not delineated by human segmentor but labeled by model. Green numbers indicate the unique cell identifier for the image.

Notably, without any immunocytochemical images in the training set segment2p also does a satisfactory job segmenting and labeling cells from these kinds of florescent images.

Note that segment2p manages to segment different size cell bodies between the two panels of Figure 2 in spite of different scales, contrast and noise levels as well as never having been trained on immunocytochemical images. Here, some post-processing of segmentation masks was necessary to compensate for the different scales – we eliminated all ROIs that included less than 50 pixels for Figure 2A and 200 pixels for 2B.

**Figure 2:**
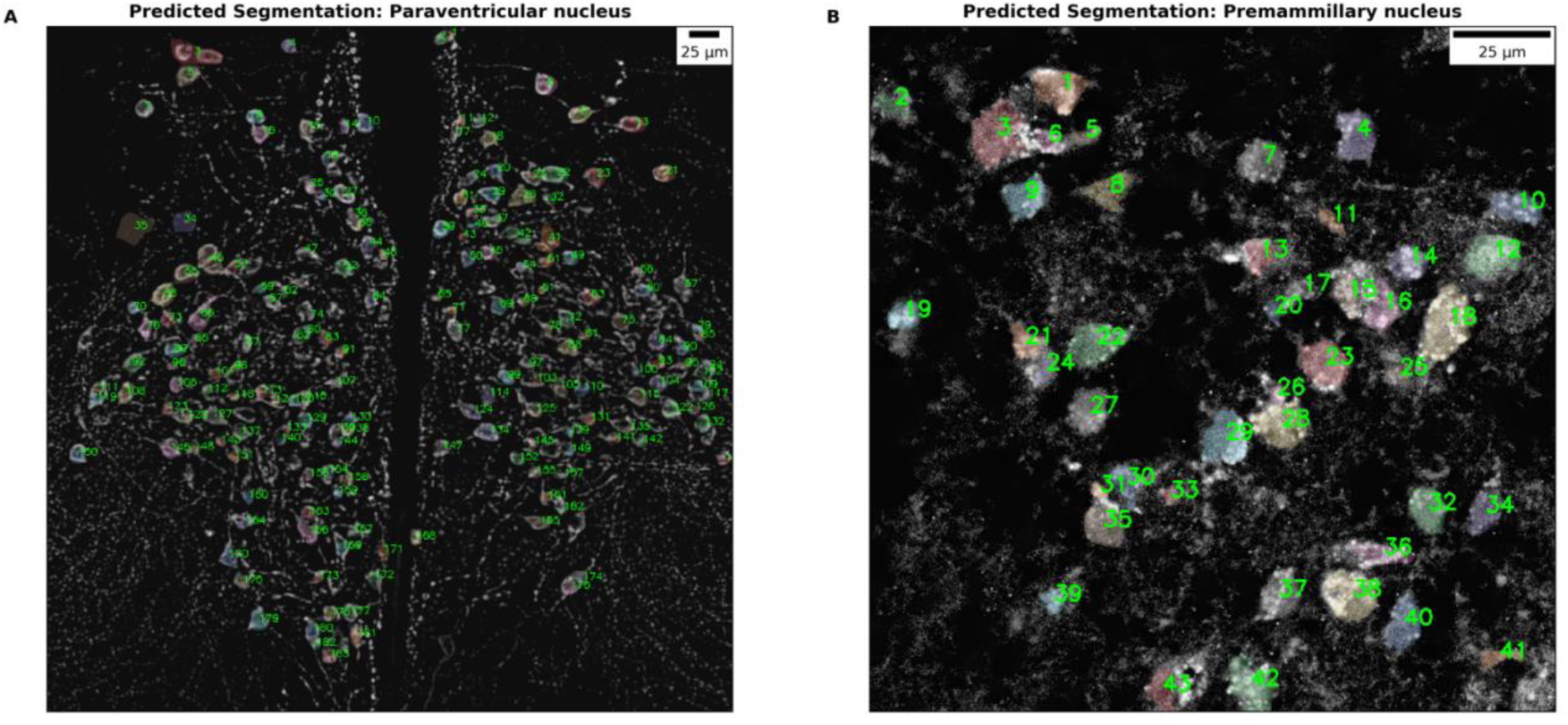
Sample segmentations of novel immunocytochemical projections. Immunocytochemical stainings overlaid with segmentation masks. (A) 30μm slice of the paraventricular nucleus of a mouse hypothalamus stained with antibodies for oxytocin. (B) Slice from the premammillary nucleus of a mouse hypothalamus stained with antibodies for histidine decarboxylase enzyme.

We designed a simple drag and drop webserver for segmenting calcium imaging micrographs. When an upload completes, the first step in the algorithm is to produce several pre-processed versions of the same file.

**Table 2:** Pre-process augmentation

| Title | Description | Designation |
| --- | --- | --- |
| Resized | Resized such that largest edge is equal to 1024 pixels and then padded such that second edge is 1024 pixels. | __orig |
| Contrast | A gamma correction is performed followed by a bilateral filter | _orig_pp |
| CLAHE | A contrast limited adjusted histogram equalization is applied | __orig_clahe |
| Distorted | After resizing such that largest edge is 1024 in size, the image is resized to a square where the sides are 90% of the largest edge and then padded to 1024. | __orig_distorted |
| Distorted Contrast | After distortion the gamma correction and bilateral filter are applied. | __orig_distorted_pp |

We include image distortion in the list of pre-processing in order to manage 2p micrographs with mixed scales in X and Y axes. In these cases, cells tend to be elongated and look very different from the original dataset (Fig 1A, Fig 3). Distortion into a square image upon submission led to satisfactory segmentation. The results are always returned in the original shape of the submission.

**Figure 3:**
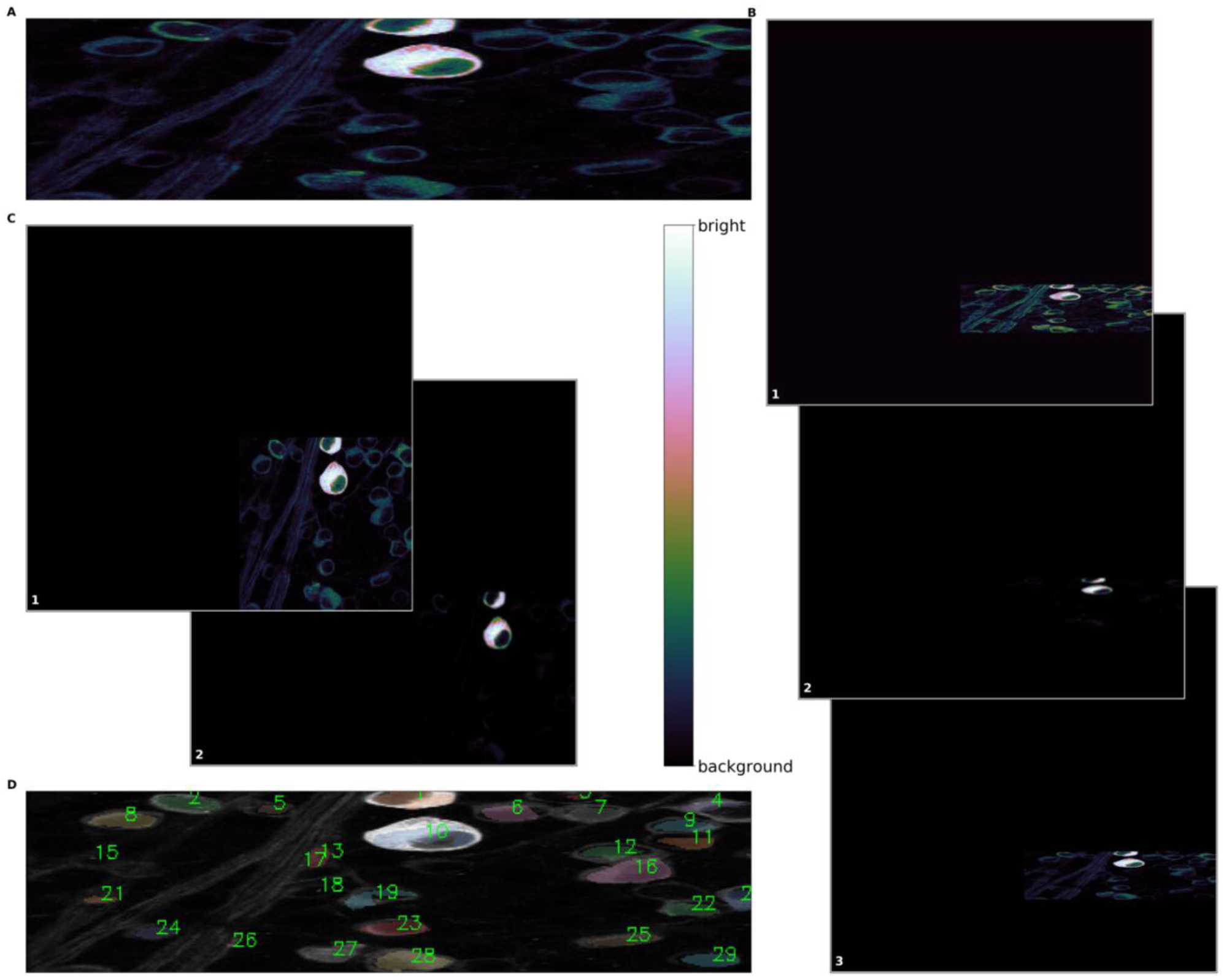
Pre-processing of input images. False color representation of a submitted calcium imaging mean projection. (A) The original image submitted. (B) Pre-processed images left undistorted. Image1, CLAHE pre-processed input image. Image 2, gamma-corrected and bilaterally filtered pre-processed input image. Image3, original image padded to be 1024 x 1024 pixels (C) pre-processed images distorted into a square. Image1, CLAHE pre-processed input image. Image 2, gamma-corrected and bilaterally filtered. (D) Resultant segmentation overlaid on original image. The algorithm converts the image in (A) to the images found in B and C (total of 5 images), segments each of them and then combines the segmentations producing the output in in D.

Not all pre-processing steps are conducive to good segmentation for every image – rather, the combination we determined is optimal for achieving reliable results with the different images we attempted to segment. We use model averaging to increase the resilience to noise and image types. Thus, after creation, the image-sets are fed into model 1 and model 2 via Amazon’s batch transform application programming interface (API). Then, the resulting binary segmentation masks are collected, the distorted masks are warped back into the original image’s shape and the masks are combined. A pixel is considered to belong to an ROI when it appears in at least two resultant masks. Since DeepLabv3 does semantic segmentation rather than instance, we separate and label neighboring ROIs using a simple erode and then watershed [11]. This method could be further improved upon by training a small convolutional network to convert the mask into instance segmentation where nearby or overlapping ROIs would likely have better separation but this would necessitate adding bounding boxes to the training set which is labor intensive. In our hands, the simple method achieves satisfactory results (cell soma detection precision ~ 70%, recall ~ 89%). The final output mask is generated such that each pixel predicted to belong to an ROI is assigned the same unique value (greater than zero) thus each ROI is assigned a distinct number label.

## 3. Validating Segment2p

In order to assess the accuracy of the algorithm, we define pixels belonging to a region of interest as one and all others as zero. The model predicts whether a particular pixel in a micrograph belongs to a region of interest. When compared to human manually segmented ground truth, a successful prediction is called a true positive. When the model predicts that a pixel belongs to a region of interest even though it does not, we call this a false positive. When the model fails to predict that a pixel is a member of an ROI even though it is, this is called a false negative. The set of pixels which are both part of an ROI in ground truth and predicted by the model we call an "interesection" (e.g., logical and). The set of pixels that are either in the ground truth or in the prediction that are part of an ROI (e.g., have a value of 1) we define as a "union" (e.g., logical or). Then we define the accuracy metric as the intersection over the union (IoU). Some of the images in our dataset were randomly left out of training to serve as a validation set for the model. The model had a validation *m*ean IoU (*m*IoU) of 0.68 +/− 0.04. This value can be affected by cells that were missed by the algorithm (false negative) or non-existing cells that were wrongly found (false positive), or because of “true positive” cells that were not identified (or not identified in their entirety) in the human segmentation.

The strength of our algorithm is its robustness to noise and image types. To quantify this, we calculate the mIoU as a function of SNR. We calculate the SNR as follows:

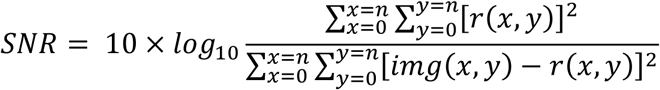

Where r is the noise image which we generate by randomly re-sampling, with replacement, pixel values outside of an ROI and n is the dimension of the image.

To quantify the accuracy as a measure of the clarity of the image, we compare signal (pixels within ROIs) to the root-mean-squared (RMS) values of the pixels outside the regions of interest from the same image. Thus, we use the following definition for SNR_RMS_ [15]:

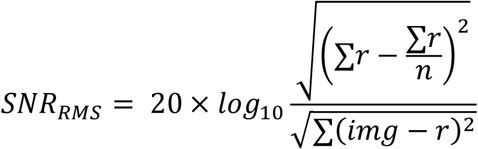

**Figure 4:**
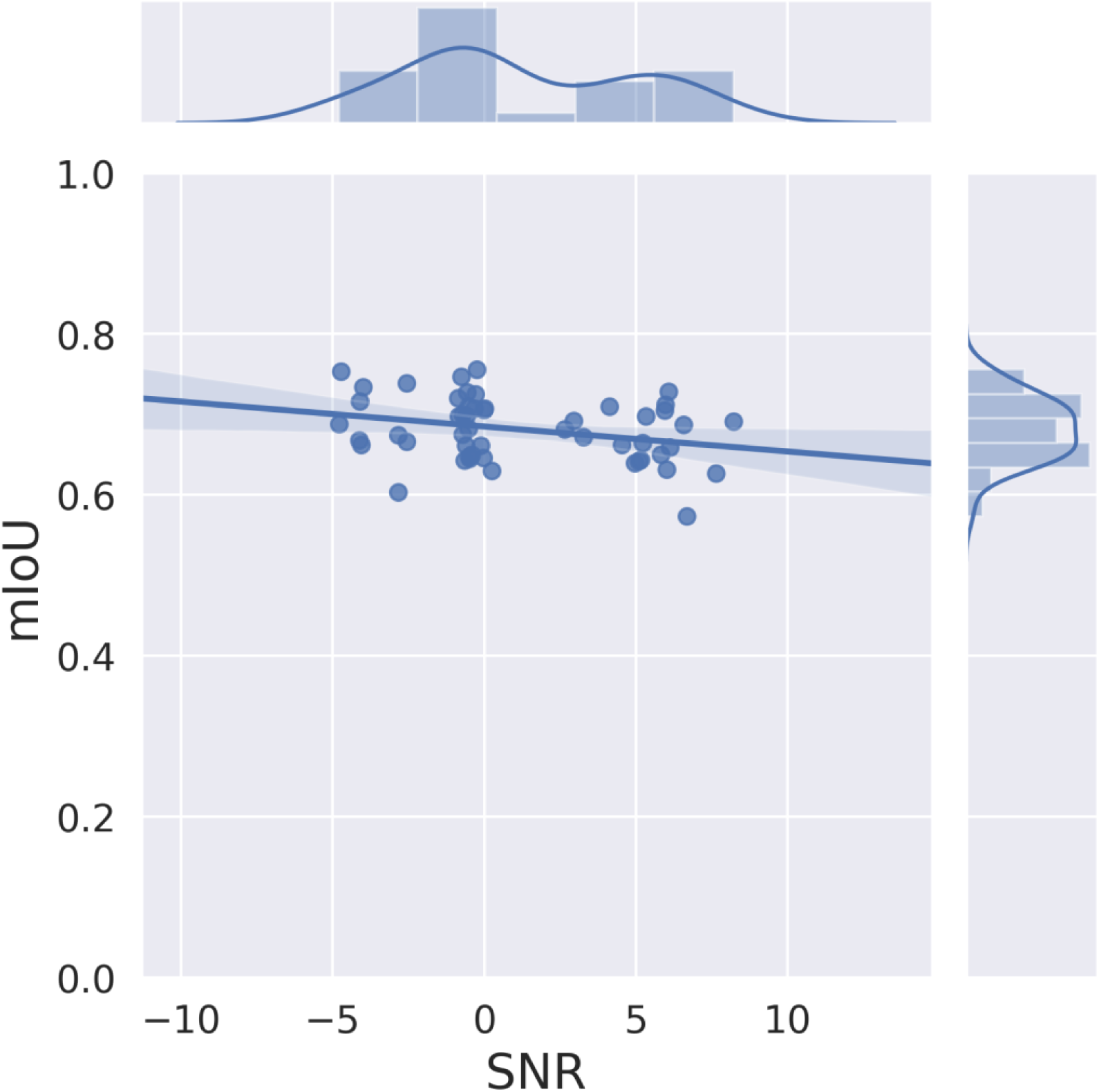
SNR vs. mIoU. The logarthmic SNR becomes more positive as the noise increases relative to the signal. The slope is estimated at −0.496 with a 95% confidence interval of [-0.502,-0.491]. Thus, a change in one decibel of SNR costs less than half an mIoU.

Then, for each image in our validation set, we calculate the SNR_RMS_and the achieved mIoU for that image. We find that the mIoU is only very slightly negative with increasing noise across our images. The slope of linear regressing SNR vs. mIoU is −0.95 × 10_−3_ CI_95%_ [−1.02 × 10_−3_, −0.88 × 10_−3_]. Thus, segment2p manages to segment well images whose ROI pixel intensities are proximal to the pixel intensities outside of ROIs, e.g., poor contrast, high background.

## 4. Discussion

Segment2p requires no installations or expertise to apply. A user can simply drag and drop a fluorescent micrograph to the website and receive an email with a segmentation mask when the process completes. To handle the submission of images to segment we designed a simple webserver which employs Dash [12] which is an open-source software for building web-based analytic applications; Redis which handles distributed data structures in memory [10] and Celery, an asynchronous job management software [9] to handle the backend preprocessing and the submission of segmentation jobs. When file uploads complete, a task is initiated and each file in the roster generates 5 additional modified submissions to both model 1 and model 2.

In addition, users can improve results simply by performing any preprocessing they deem appropriate and submitting these files along with the originals. For instance, by running a contrast correction or performing motion correction prior to taking the mean/max projection. The algorithm will automatically combine the segmentations from these preprocessed submissions whereby a pixel is considered belonging to an ROI when it appears at least twice. In case additional improvement is required for a particular submission, the public GitHub repository [8] can be cloned easily (see the readme in the repository for instructions). The model can then be fine-tuned on new data.

Due to the large number of cells in our calcium image micrographs, often the ground truth as established by human segmentors, miss real cells that the algorithm catches [Fig 1 c-d]. In addition, in the presence of neuropil, there is not 100% agreement between expert human segmentors, on what constitutes a cell soma and thus we propose that the SNR of the extracted calcium traces be used as a guide to eliminate ROIs which do not show calcium activity (not provided). With some modification, these segmentations should also be able to serve as seeds for the CNMFe algorithm [2, 3].

Converting this algorithm to a real-time model is feasible since it takes each model about 1/10_th_ of a second to process an image on an Amazon p3.2xlarge instance. Further speed up can be achieved by implementing a Sagemaker pipeline for file ingestion. To implement a real-time model, one solution would be to perform a baseline recording, collect several calcium imaging micrographs, average and pre-process them then submit them to segment2p for segmentation. Such a further development would enable feedback experiments or stimulations conditional on observed neural activity.

## 5. Conclusion

A set of mean or max projections from a calcium imaging stacks are submitted for segmentation via our website, www.segment2p.co.il. After adding an e-mail address and clicking submit an e-mail is sent to the inputted address.

**Figure 5:**
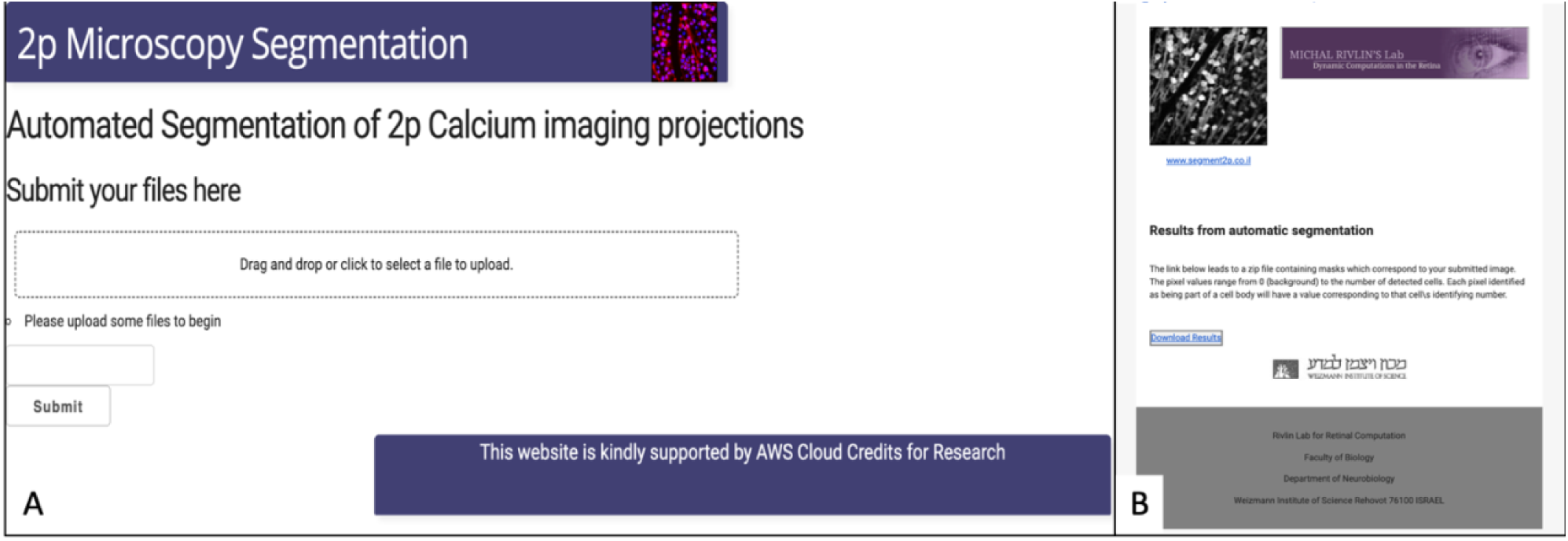
Website and results email snapshot. (A) A snapshot of our online submission system www.segment2p.co.il. After uploading images and writing an email address, clicking submit will launch the segmentation job. (B) Results are received via e-mail with a link to a zip file containing the masks corresponding to the submitted images.

Behind the scenes, the image is cloned and pre-processed and then submitted to two segmentation networks. The results are combined and a merged mask is provided. In our hands, we found that different brain regions, 2p microscopy and immunocytochemical stains are all well segmented by the algorithm.

Currently, segment2p is freely available both as a web service and the source code is in a public Github repository. In addition, the training dataset and model weights are available upon request. Segment2p is a reliable solution for segmenting fluorescent micrographs being both easy to use as well as robust to noise, cell types and imaging techniques.

## 6. Acknowledgments

We would like to thank Amazon Web Services for their generous research credit award. We’d also like to thank the Kimchi lab for being early beta testers and for providing the data for figure 3A. This project was supported by research grants from the Israeli Centers Of Research Excellence (I-CORE) (51/11), the Minerva foundation, the Israel Science Foundation (ISF) foundation (1396/15) and the European Research Council (ERC) under the European Union’s Horizon 2020 research and innovation program (grant agreement No. 757732).

